# Dynamic Reworking of Marine Diatom Endometabolomes

**DOI:** 10.1101/2025.07.25.666871

**Authors:** Malin Olofsson, Mario Uchimiya, Frank X. Ferrer-González, Jeremy E. Schreier, McKenzie A. Powers, Christa B. Smith, Arthur S. Edison, Mary Ann Moran

**Affiliations:** Department of Marine Sciences, University of Georgia, Athens, GA 30602, USA; Department of Aquatic Sciences and Assessment, Swedish University of Agricultural Sciences, Box 7050, 750 07 Uppsala, Sweden; Complex Carbohydrate Research Center, University of Georgia, Athens, GA 30602, USA; Department of Biochemistry and Molecular Biology, University of Georgia, Athens, GA 30602, USA

**Keywords:** diatoms, bacteria, endometabolites, temperature acclimation, co-culture Running title: Diatom endometabolome dynamics

## Abstract

A large annual carbon flux occurs through the ocean’s labile dissolved organic carbon (DOC) pool, with carbon influx dominated by phytoplankton-derived metabolites and outflux by heterotrophic bacterioplankton uptake. We addressed the dynamics of this flux between marine primary and secondary producers through analysis of the *Thalassiosira pseudonana* CCMP1335 endometabolome, a proxy for labile DOC release during phytoplankton excretion and mortality. Diatom strains acclimated at one of three different temperatures (14°C, 20°C, or 28°C) were then cultured either axenically or with the bacterium *Ruegeria pomeroyi* DSS-3, and their endometabolites analyzed by NMR. Osmolytes were by far the most dynamic, exhibiting concentration differences up to 150-fold between conditions; median concentration variation across identified endometabolites was ∼1.5-fold. Differential expression of diatom metabolic pathways suggested changes in synthesis rates as a mechanism for endometabolome remodeling. Consistent with expectations of high turnover, endometabolite mean lifetimes prior to bacterial uptake were <2 h to 12 h.

**Importance:** The role of labile DOC in the transfer of marine carbon between phytoplankton and heterotrophic bacteria was first recognized 40 years ago, yet the identity and dynamics of phytoplankton metabolites entering the labile DOC pool are still poorly known. Using metabolome and transcriptome profiling, we found dynamic composition and concentration of diatom endometabolites, depending on growth conditions and arising over time frames as short as a single growth cycle. This strong response to external conditions, both biotic and abiotic, has implications for downstream processing and fate of ocean carbon by heterotrophic bacteria.

## Introduction

In the microbe-metabolite network of the surface ocean, a large fraction of heterotrophic bacterial production is supported by the organic carbon released by marine phytoplankton into the dissolved organic carbon (DOC) pool (1, 2). The mechanisms by which labile metabolites enter this carbon reservoir fall into three categories: release from living phytoplankton (both diffusive and active processes); release from phytoplankton cell rupture (zooplankton grazing, viral lysis, and senescence) (3–5); and release from heterotrophs (secondary producer exudates and waste) (6). The first two of these mechanisms liberates a mixed suite of internal metabolites from phytoplankton cells that can include polysaccharides (7), organic sulfur compounds (8), sugars, amino acids (2), and osmolytes (9). Quantitatively, phytoplankton endometabolite release by excretion and rupture mechanisms is estimated to account for ∼80% of labile DOC inputs to surface seawater (1).

Once metabolites are released into the DOC pool, their uptake and processing by bacteria represents a major carbon flux at the global scale. In model organism studies (10, 11) and natural bacterial communities (12), marine bacteria metabolize components of the mixed metabolite pool selectivity, sometimes referred to as ’substrate preference’, likely driven by genomic constraints, enzyme kinetic properties, or regulatory processes. At a broad taxonomic level, members of the *Roseobacteraceae* take up a range of organic acids and sulfur-containing metabolites (13, 14), whereas *Flavobacteriaceae* dominate polysaccharide degradation (10, 13, 15), and *Gammaproteobacteria* target peptides, polysaccharides, and monosaccharides (12). Some studies suggest substrate energy content as a driver of preferential bacterial uptake (16), while others propose a metabolic specialization predictable from genome content (11, 17). Regardless of the underlying mechanism, the chemical composition of intracellular pools released by phytoplankton is likely to impact downstream processing by heterotrophic bacterioplankton communities.

As ocean temperatures shift on annual and decadal scales (18), the consequences to phytoplankton endometabolome composition are hard to anticipate. Here we ask whether acclimation of diatom strains at different temperatures affects their accumulation of internal metabolites, and secondarily whether there are interaction effects on endometabolome composition when heterotrophic bacteria are present. From previous research, temperature shifts have been shown to alter phytoplankton physiology by changes in cell size (19, 20), shifts in chlorophyll *a* concentration (21, 22), and alterations of protein content (21). Further, increased temperature may impact the mechanisms that release endometabolites by cell breakage during mortality, including timing of viral infections in phytoplankton blooms (23) and rates of zooplankton feeding (24). If temperature also affects endometabolite concentration or composition, resultant changes to the labile DOC pool could have repercussions for bacterial carbon processing.

In this study, the marine diatom *Thalassiosira pseudonana* CCMP1335 was acclimated over three months at three temperatures, one at the diatom’s optimal growth temperature (20°C) (22), one below (14°C), and one above (28°C). Following acclimation, diatom co-cultures were inoculated with the marine bacterium *Ruegeria pomeroyi* DSS-3 or were left axenic. Diatom endometabolites were examined by three methods: identification by nuclear magnetic resonance (NMR) spectroscopy, analysis of expression of metabolite biosynthesis pathways, and assessment of lability via bacterial drawdown assays. The convergence of these datasets provides evidence of considerable temperature-related restructuring of diatom endometabolomes, points to a substantial impact of heterotrophic bacteria on endometabolome composition, and singles out osmolytes as the organic compound class most responsive to conditions in the diatom’s environment.

## Results and Discussion

### Axenic *T. pseudonana* endometabolomes are restructured during temperature acclimation

Axenic cultures of the diatom *T. pseudonana* CCMP1335 were pre-acclimated at 14°C, 20°C, and 28°C (Fig. 1a) under replete nutrient and vitamin conditions over three months with weekly transfers (∼120 generations). Following acclimation, an experiment was established to compare endometabolome composition across the temperatures and in the presence or absence of a heterotrophic bacterium (Fig. 1b). These cultures were incubated over one growth cycle at their temperature of acclimation until late exponential phase, then harvested and analyzed for endometabolome composition.

**Fig. 1.**
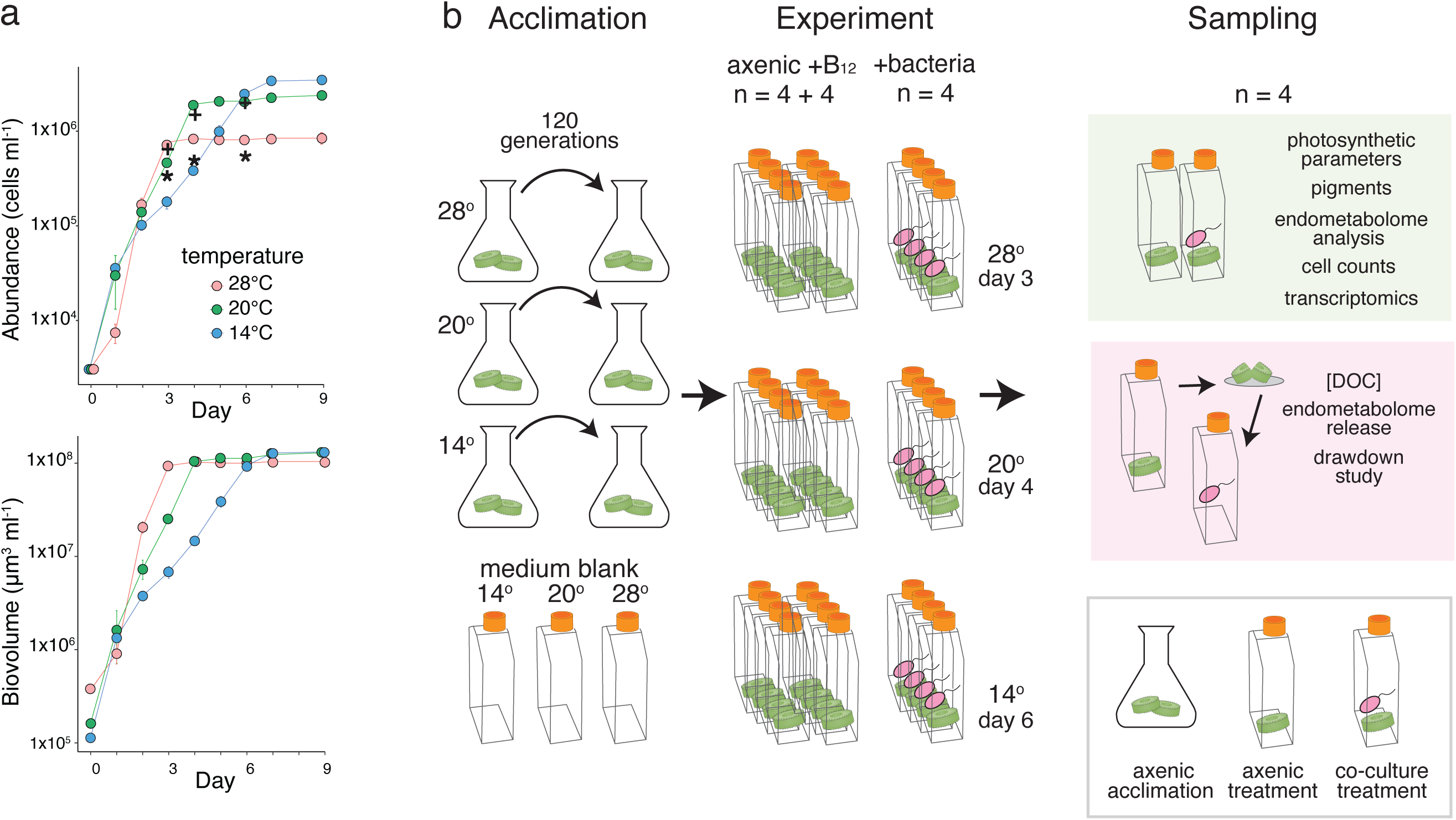
(a) *T. pseudonana* cell abundance (top) and biovolume (bottom) in pre-experiment axenic cultures following temperature acclimation for 120 generations were used to determine the timing of late exponential phase (n = 2). Error bars indicate standard deviation; most fall within the symbols. In the top panel, *T. pseudonana* cell abundance in the main experiment is also indicated, with * (co-cultures) and + (axenic cultures) symbols indicating cell number on the day of harvest (28°C cultures on day 3; 20°C cultures on day 4; 14°C cultures on day 6). (b) In the main experiment, temperature acclimated *T. pseudonana* cultures were inoculated with *R. pomeroyi* or remained axenic. At harvest, phytoplankton cells were collected for physiological measurements, endometabolome analysis, and transcriptomics (green shading). In a separate metabolite drawdown study (pink shading), axenic diatom endometabolomes were inoculated with *R. pomeroyi* to measure lability. DOC, dissolved organic carbon.

Because exponential growth was offset in time across the temperatures (Fig. 1a), consistent with expectations in nutrient replete conditions (25), cultures were harvested at 3 d (28°C), 4 d (20°C), or 6 d (14°C). Cell numbers were higher at 14°C but cell volumes were higher at 28°C (Table S1), the latter in accordance with data on temperature influence on *T. pseudonana* cell size (22, 26) and consistent with reports of diatoms being an exception to the negative size-temperature relationship (20, 27, 28). Because of cell size differences, diatom biovolume was similar at stationary phase (Fig. 1a). Photochemical efficiency measurements were indicative of healthy cells (Fv/Fm values >0.70; Table S1).

Endometabolomes were liberated from the axenic diatom cultures at harvest to characterize internal metabolites, serving as a proxies for compounds released by excretion or mortality processes. Sixteen metabolites were annotated by NMR (Fig. 1b, Table S2). Ten had significantly different concentrations for at least one of the three pairwise temperature comparisons (14°C vs. 20°C, 14°C vs. 28°C, and 20°C vs. 28°C) (Kruskal Wallis with post-hoc pairwise Mann-Whitney U tests, *P* ≤ 0.05) (Fig. 2a). Of these, six had directional changes in which concentration changed in tandem with temperature (i.e., significantly increasing or decreasing along the temperature gradient) or in a threshold pattern (one significant change along the temperature gradient).

**Fig. 2.**
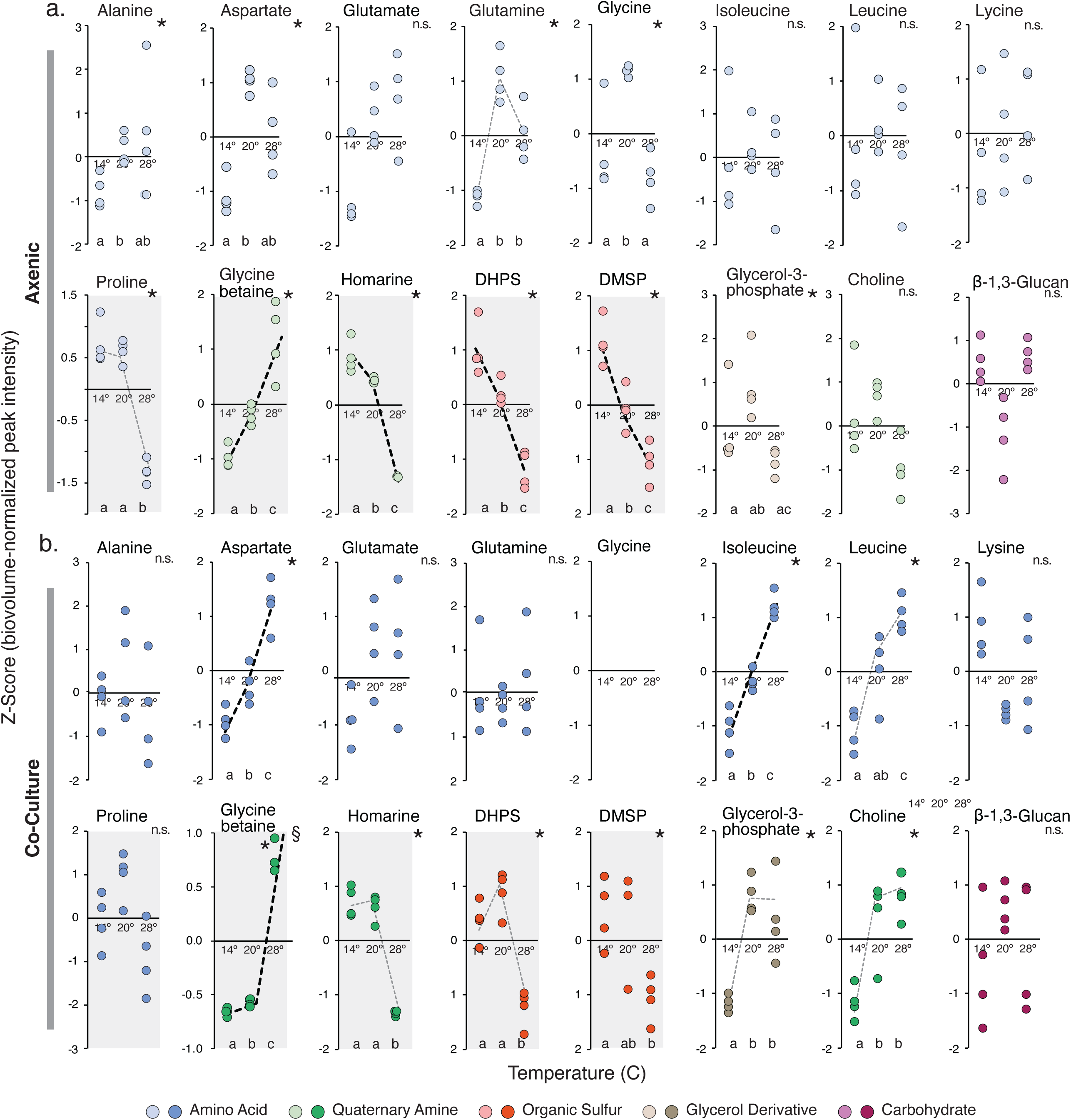
Abundance of endometabolites in temperature acclimated *T. pseudonana* strains. Peak intensity is normalized to diatom biovolume at time of harvest and presented as Z-scores for (a) axenic cultures (light symbols) and (b) co-cultures (dark symbols). Asterisks indicate differences in metabolite concentrations that are significant at P ≤ 0.05 (Kruskal Wallis Test). Dotted lines highlight metabolites that trend with temperature as either a tandem (bold black lines) or threshold (light gray lines) response. Osmolytes are indicated by gray-shaded plots. Unalike lower-case letters at the bottom of each plot indicate significant concentration differences between acclimation temperatures in post-hoc pairwise comparisons at P ≤ 0.05 (Mann-Whitney U Tests). n.s., no significant differences. §, one replicate for glycine betaine at 28°C plots above the Y axis at 2.4 and is not shown.

Considering both of these patterns, glycine betaine and glutamine concentrations were positively related to temperature, and proline, homarine, 2,3-dihydroxypropane-1-sulfonate (DHPS), and dimethylsulfoniopropionate (DMSP) concentrations were negatively related (Fig. 2a). Many of the temperature responsive metabolites function as osmolytes (e.g., proline, glycine betaine, homarine, DHPS, and DMSP) (9), and typically have high concentrations in marine phytoplankton cytosols. For example, previous measures of these endometabolites in axenic *T. pseudonana* cells ranged from 16 mM to 118 mM (8, 29, 30). Named originally for their role in regulating osmosis based on changes in external water potential (31), osmolytes have also been shown to alleviate temperature stress by inhibiting protein denaturation (32). Indeed, osmolyte concentration changes in phytoplankton endometabolomes in response to temperature have been reported previously in both laboratory and field studies (30, 33, 34).

Nine amino acids were also among the axenic *T. pseudonana* endometabolites, five of which appeared responsive to acclimation temperature based on differences in concentration for at least one of the three pairwise comparisons. Concentrations of two amino acids tracked with temperature, including proline, both an osmolyte and an amino acid, and glycine (Fig. 2a) (22). Overall, compounds dominating the diatom endometabolomes overlapped with those observed in natural phytoplankton communities. For example, in a North Pacific Ocean study, glycine betaine, and proline were among the most abundant osmolytes; and alanine, aspartate, glutamate, glutamine, and leucine were among the most abundant amino acids (33).

### *T. pseudonana* endometabolomes differ between co-culture and axenic conditions

The *T. pseudonana* endometabolome composition was also analyzed after acclimated strains were grown in co-culture with a heterotrophic marine bacterium (Fig. 2b), as would occur in a natural seawater environment. Cultures to be inoculated with *Ruegeria pomeroyi* DSS-3, a representative of the *Roseobacteraceae* family whose members are frequently found in association with phytoplankton blooms (12), were first stepwise limited by B_12_, a vitamin that is supplied exogenously by associated marine bacteria in nature. Thus *R. pomeroyi* was the only source of B_12_ for the diatom, and diatom-derived DOC was the only substrate for the bacterium. All co-cultures were incubated at their temperature of acclimation for one growth cycle until harvest, as for the axenic cultures. The co-cultured diatoms’ growth rates were slower than in axenic cultures (Fig. 1a, Table S1) and suspected to be related to the bacterial B_12_ supply rate. This was supported by the enrichment of the diatom’s B_12_ acquisition protein CBA1 (35) in co-culture transcriptomes compared to their axenic counterparts, with 5.7 log_2_-fold differences in relative gene expression averaged across the three acclimation temperatures (Tables S3-S5).

Only fifteen compounds were identified in the co-cultured diatom endometabolomes because glycine was not detected. Three had temperature responses that followed the patterns of the axenic cultures; these were osmolytes glycine betaine (increased concentration with increasing temperature), homarine, and DHPS (decreased concentration with increasing temperature). Five others had significant temperature responses that were not observed under axenic conditions; these were three amino acids (aspartate, isoleucine, and leucine), glycerol-3-phosphate, and choline, which all increased in concentration with increasing temperature (Fig. 2b).

While trends of endometabolite changes with acclimation temperature were largely consistent between the axenic and co-culture diatoms, primarily for the osmolytes (Fig. 2), the metabolite concentrations differed substantially. Twelve of the 16 identified metabolites had statistically different concentrations in co-culture versus axenic treatments for at least one acclimation temperature (Wilcoxon Rank Sum Test, Bonferroni multiple tests correction, *P* ≤ 0.05; Fig. 3). Three had higher concentrations in the co-cultures compared to axenic: β-1,3-glucan (4-fold, averaged means of the three temperatures), lysine (4-fold), and proline (2-fold). Nine had lower concentrations in co-cultures, with the most extreme differences found for glycine betaine (97-fold), DMSP (16-fold), and glycine (not detected in co-culture endometabolomes) (Fig. 3). A further comparison of the full suite of peaks in the NMR spectra (i.e., whether or not they were assigned to a known compound) similarly showed that interfacing with the bacterium for one growth cycle altered concentrations of the majority of diatom endometabolites (Fig. S1). These data portray diatom endometabolomes as highly dynamic in composition and decidedly responsive to environmental conditions linked to both temperature changes and interactions with bacteria. We explored a possible temperature acclimation mechanism with transcriptome data from the *T. pseudonana* strains.

**Fig. 3.**
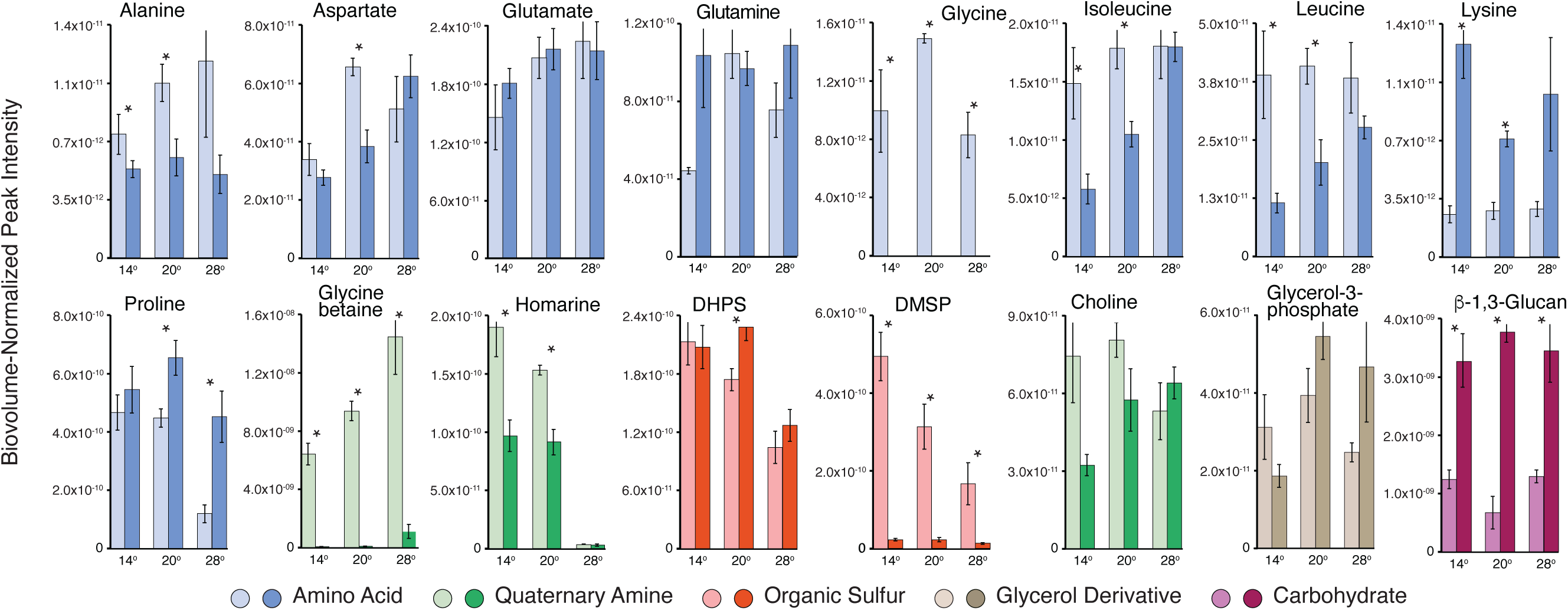
Comparison of endometabolome concentrations in temperature-acclimated *T. pseudonana* strains from axenic cultures (light bars) and co-cultures (dark bars). Peak intensity is normalized to diatom biovolume at time of harvest. Error bars show ±1 standard deviation, n = 4. Asterisks above each co-culture/axenic pair denote differences in concentration based on Student’s T-tests with Benjamini-Hochberg multiple comparison corrections (n = 4, P ≤ 0.05).

### *T. pseudonana* metabolite biosynthesis pathways have temperature-specific expression patterns

The changes in the diatom endometabolome pools in response to variations in environmental conditions, regardless of ultimate mechanism, must result from a proximal effect on input rates (i.e., metabolite biosynthesis) and/or on output rates (e.g., metabolites repurposed internally or released externally). To address the former, transcriptomes of co-cultured *T. pseudonana* were compared among the temperature-acclimated strains. Of the 11,675 predicted genes, ∼7,000 had significantly altered relative expression between the 14°C and 28°C transcriptomes (Table S7). For fourteen metabolite biosynthesis pathways tentatively identified in the genome (Table S6), five had more genes enriched at 14°C (up to 30-fold; DESeq2, *P*adj ≤ 0.05), and eight had more genes enriched at 28°C (up to 2-fold; DESeq2, *P*adj ≤ 0.05) (Fig. 4). The pairwise comparisons with the 20°C transcriptome were largely intermediate to these results for these temperature end points.

**Fig. 4.**
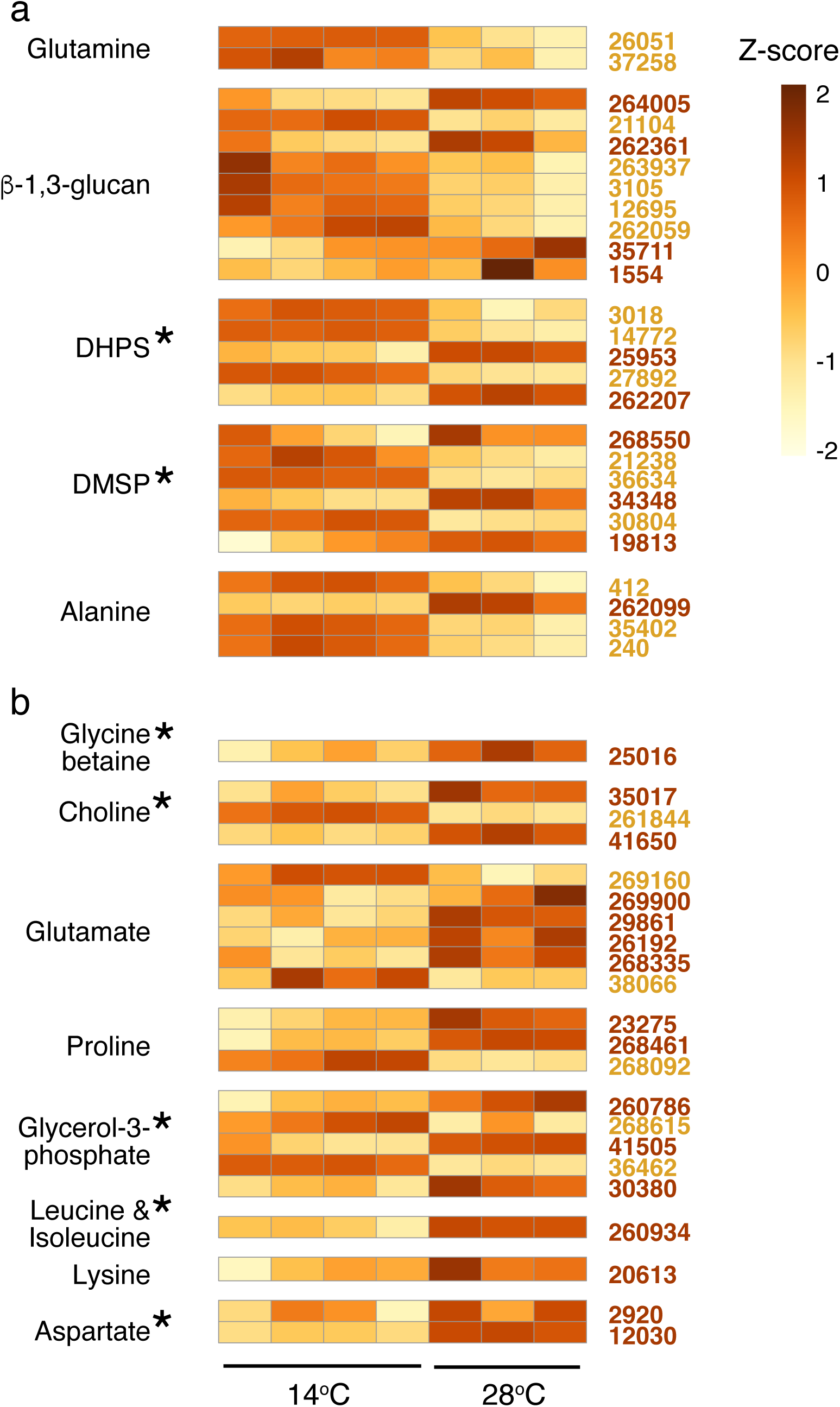
*T. pseudonana* gene expression of endometabolite biosynthetic pathways with reactions biased toward enriched expression in 14°C transcriptomes or 28°C transcriptomes, normalized as Z-scores. Each row represents a gene in a tentative biosynthetic pathway and each column represents a treatment replicate. (a) Pathways with the majority of significant genes enriched at 14°C (gene locus tags in gold font). (b) Pathways with the majority of significant genes enriched at 28°C (gene locus tags in brown font). Asterisks indicate significant differences in concentration between 14°C and 28°C as shown in Fig. 2 that are consistent with pathway expression. One replicate of the 28°C-acclimated cultures was an outlier and is not included.

There was good agreement between the direction of expression changes in tentative metabolite synthesis pathways (Fig. 4) and the direction of concentration shifts in the diatom’s endometabolome (Fig. 2b). DHPS and DMSP had greater biosynthetic pathway enrichment and higher endometabolome concentrations at 14°C relative to 28°C. Aspartate, leucine, isoleucine, glycine betaine, glycerol-3-phosphate, and choline had higher biosynthetic pathway enrichment and higher concentrations at 28°C relative to 14°C. These matches between metabolite concentrations and directional change in gene expression suggest regulatory changes in synthesis pathways as a mechanism of temperature acclimation. Other hints of cellular acclimation mechanisms from the transcriptomes include changes in metabolite exudation from photosynthetic overflow or photorespiration (5), consistent with significant enrichment of transcripts for two central photorespiration genes: phosphoglycolate phosphatase (1.4-fold) and (S)-2-hydroxy-acid oxidase (1.4-fold) in the 28°C acclimated strain; and changes in metabolite concentration for stress reduction, evidenced as enrichment in catalase and superoxide dismutase transcripts (14°C strain; 1.3-to 1.8-fold) and in nutrient acquisition, evidenced as enrichment of four of the five ammonium transporters (28°C transcriptome; 1.3-to 1.9-fold) (Table S7).

### Bacterial drawdown rates indicate lability of *T. pseudonana* endometabolites

*R. pomeroyi* abundance increased an average of 5.6-fold (± 0.8) in *T. pseudonana* co-cultures between inoculation and late exponential phase harvest. As phytoplankton endometabolites were the only source of substrates available to the bacterium, we conducted drawdown experiments to determine which of the identified metabolites were supporting *R. pomeroyi* growth. Cell lysates collected from a second set of axenic flasks (n = 4 for each acclimation temperature; Fig. 1b) were diluted to 204 ± 12 µM C, brought to 30°C, and inoculated with *R. pomeroyi*. Concentrations quantified by NMR analysis at the time of inoculation and again after 10 h indicated concentration decreases in all endometabolites except one: β-1,3-glucan, a component of the storage molecule laminarin (Fig. 5). This metabolite can be readily degraded by other marine bacteria (12, 36), but the *R. pomeroyi* genome lacks a laminarinase gene.

**Fig. 5.**
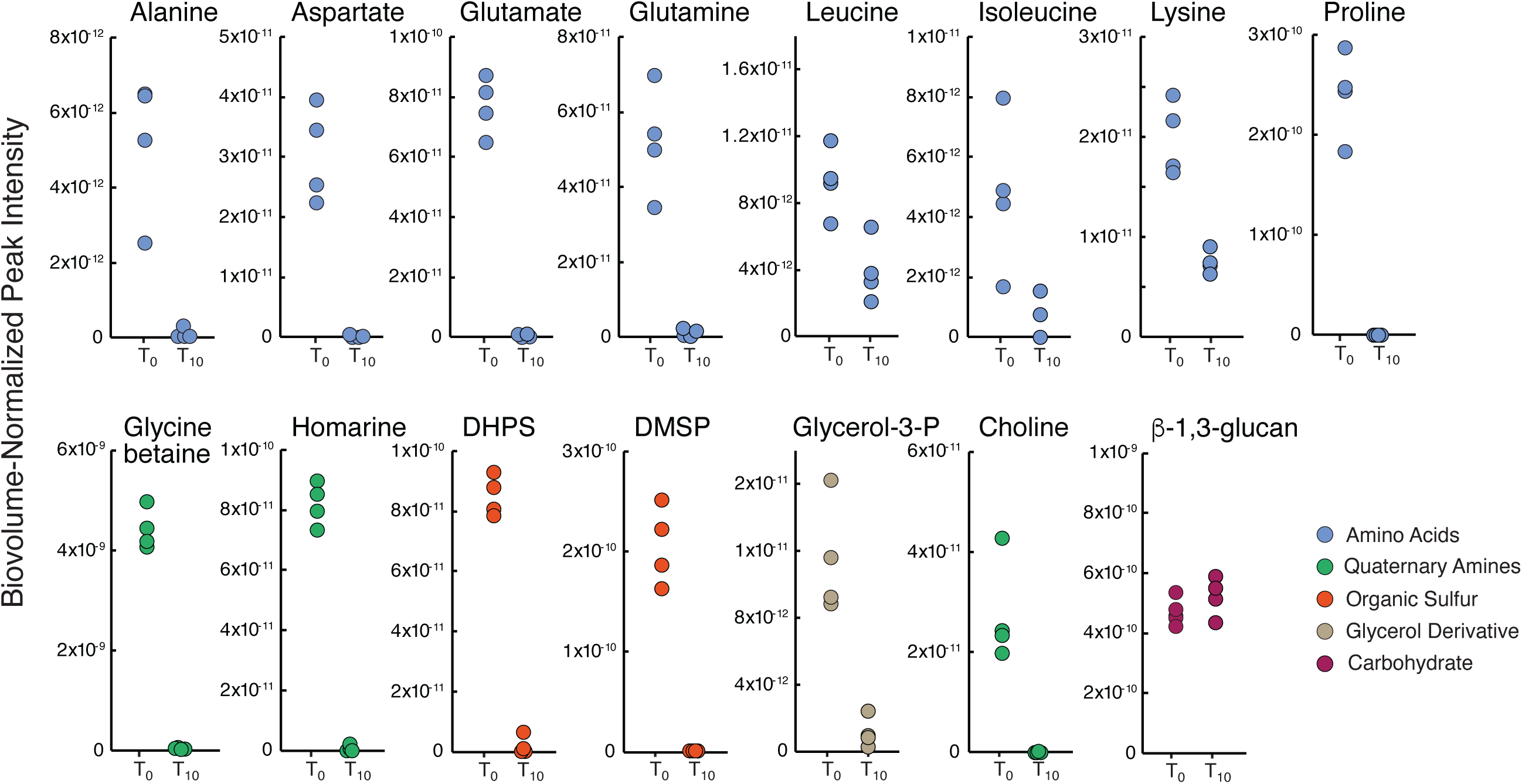
Endometabolite drawdown experiment measuring *R. pomeroyi* assimilation of compounds in cell lysates from temperature-acclimated axenic diatoms. Metabolite concentrations were measured at the time of inoculation (T_0_) and after 10 h (T_10_). Data are from the 20°C-acclimated diatom treatment; drawdown results at the other two temperatures are consistent (Fig. S2). Endometabolite peak intensity is normalized to diatom biovolume at the time of harvest. Concentrations at inoculation compared to 10 h are significantly different for all metabolites except β-1,3-glucan (Student’s T-test with Benjamini-Hochberg correction, P ≤ 0.05; n = 4).

For 10 metabolites, final concentrations were ≤ 2% of initial. Thus alanine, aspartate, glutamate, glutamine, proline, glycine betaine, homarine, DHPS, DMSP, and choline had mean lifetimes of 2 h or less in the substrate pool. The rapid bacterial metabolism of these diatom-derived compounds is consistent with earlier findings that they represent major currencies of surface ocean carbon flux (10, 37–39). Metabolites isoleucine, glycerol-3-phosphate, lysine, and leucine had mean lifetimes of 4, 5, 10, and 12 h, respectively, with drawdown to concentrations between 90% and 60% of initial. Metabolites in this second group are less likely to support *R. pomeroyi* growth when other bacterial species are competing for access to substrates, reiterating the importance of the chemical makeup of the labile DOC pool in the success of bacteria associating with phytoplankton cells (11, 12). For example, isoleucine and leucine were among the more slowly assimilated metabolites (Fig. 5) and also among the compounds that increase in concentration with *T. pseudonana* acclimation temperature (Fig. 2b). Thus an overlay of species-specific bacterial uptake rates onto endometabolome composition may have predictive value for bacterial community composition shifts under different temperature regimes (40).

At a common temperature of 30°C, bacteria in the drawdown experiment assimilated metabolites at the same rate regardless of acclimation temperature of the diatom strain. Thus, as expected, lability was not affected by acclimation temperature (22) (Student’s T-test, *P* ≤ 0.05; n = 3 or 4). There was one exception, however, in which DHPS was taken up more slowly from the 14°C endometabolome, being drawn down to only 30% of initial concentration compared to 1% of initial in 20°C and 28°C endometabolomes (Fig. S2). No other metabolites in the 14°C cell lysates had lower drawdown rates, ruling out an inhibitory compound in endometabolome. However, other compounds whose concentrations peaked in the 14°C endometabolomes may be a preferred substrate of the bacterium; candidate metabolites are proline, homarine, and DMSP (Fig. 2a).

### A convergence of evidence points to dynamic compositional shifts in the *T. pseudonana* endometabolome

Acclimation of axenic *T. pseudonana* at three different temperatures over ∼120 generations resulted in median endometabolite concentration differences of 1.5-fold across all pairwise comparisons of the 16 metabolites (46% were higher at the higher temperature, 54% were lower; n = 48); and co-culturing *T. pseudonana* strains with a bacterium resulted in median endometabolite differences of 2-fold (37% were lower than axenic, 63% were higher). In these comparisons, osmolytes were the most responsive when compared across growth conditions (Figs. 2,3). Such concentration dynamics have indeed been documented previously, for example for glycine betaine in natural phytoplankton endometabolomes (33) and DMSP in sea-ice diatom endometabolomes (30), and are not surprising given the role of osmolytes in maintaining salt homeostasis (9), preventing protein denaturation (32), scavenging reactive oxygen species (41), maintaining membrane fluidity and cryoprotection (32), and encouraging growth of beneficial bacteria (42, 43). Metabolites with the largest differences across all comparisons were glycine betaine and DMSP (between co-culture and axenic conditions, 167-fold and 16-fold) and homarine (between acclimation temperatures, 33-fold).

However, the most extreme concentration shift in the *T. pseudonana* endometabolomes was for glycine, which was present in axenic culture but not detectable in co-cultures (Fig. 3). The strong response of this particular metabolite to *R. pomeroyi* presence is consistent with evidence that exogenous glycine has diverse effects on bacterial physiology, including enhancement of oxidative stress (44) and the induction of morphological changes in membrane lipopolysaccharides (45). The laminarin component β-1,3-glucan, which the bacterium did not utilize, also had dynamic concentration responses in *T. pseudonana* exometabolomes which might indicate bacterially-induced alteration of carbon storage (Fig. 3). Such substantial reworking of diatom endometabolite composition based on environmental conditions is consistent with previous observations of vitamin-limitation responses (46), growth stage responses (15), and salinity responses (30).

Phytoplankton endometabolites enter the labile DOC pool through selective release by healthy cells (5), or as a mixed metabolite pool liberated by zooplankton ‘sloppy feeding’ (6, 47) or viral lysis (4). Fifteen of the 16 labile organic compounds identified here also contain nitrogen, phosphorus, and/or sulfur, playing key roles in microbial physiology and biogeochemistry. Bacterial uptake of *T. pseudonana* metabolites was both rapid and preferential (Fig. 5). Of the compounds most responsive to environmental conditions in this study, two have been proposed previously to be major sources of bacterial carbon in the surface ocean: glycine betaine, estimated to account for up to 35% of heterotrophic bacterial carbon demand (48), and DMSP estimated to account for up to 10% (49, 50). The link between the production of metabolites by phytoplankton and their consumption by bacteria, mediated through the labile DOC pool, represents a highly dynamic and poorly characterized sequence in the ocean’s biogeochemical cycles.

## Methods

### Culture conditions

Axenic cultures of the diatom *Thalassiosira pseudonana* CCMP1355 (National Center for Marine Algae) were acclimated to three temperature conditions (below optimal: 14°C, optimal: 20°C, and above optimal: 28°C) for three months with weekly transfers (∼120 generations). These temperatures were selected based on previous growth rate measurements of the diatom strain across a range of temperatures (22). Diatoms were cultured in L1 medium under nutrient-replete conditions (883 µM NO_3_, 500 µM PO_4_) (51) at a salinity of 35 (52) in acid-washed glass containers under 120 µmol photons m^-2^ s^-1^ (ULM-500 Light Meter, Walz) with a 16:8 h light:dark cycle. The starter cultures were checked for purity at the initiation of the acclimation and at each weekly transfer by plating aliquots on ½ YTSS medium (53).

For the main experiment, 39 culture flasks (1.9 L Nunc polystyrene flasks) were prepared with 1 L of modified L1 medium with NaH^13^CO_3_. *T. pseudonana* was inoculated into 36 flasks, 12 per acclimation temperature, at ∼3 x 10^4^ cells ml^-1^. For the 12 co-culture treatment flasks, B_12_ was limited through 3 weekly transfers into L1 medium with 100-fold decreased B_12_ concentration. These cultures were inoculated with alphaproteobacterium *Ruegeria pomeroyi* DSS-3, representing a bacterial taxon typically associated with phytoplankton blooms (12). Prior to inoculation, the bacterium was grown overnight in ½ YTSS medium, harvested in exponential growth phase, and washed five times in sterile artificial seawater at 6,000 RCF. Bacteria were inoculated into the 12 co-culture flasks at ∼1 x 10^5^ cells ml^-1^, with four replicates per acclimation temperature. Co-culture medium had no added vitamin B_12_ (which was instead provided by *R. pomeroyi*); axenic culture medium had B_12_ added to a final concentration of 0.37 nM (Fig. 1b). A flask containing medium but no microbes was established for each treatment as controls for metabolite analysis. The *T. pseudonana* axenic cultures were checked for purity at the initiation of the experiment by plating on ½ YTSS, and again when the experiment was terminated at late exponential phase by plating and inspection of flow cytometry data for bacterial signatures.

Flasks were harvested at late exponential growth phase on days 3 (28°C), 4 (20°C), and 6 (14°C), with sampling occurring 7 h into the light cycle (Fig. 1a, n = 2). The axenic flasks (n = 24, 600 ml) were immediately filtered onto 2.0 µm-pore-size Isopore filters (Millipore, Burlington, MA) to collect diatom cells, and filters were stored at -80°C for analysis of axenic endometabolomes (12 filters) and as metabolite sources in the drawdown experiment (12 filters). The co-culture flasks (n = 12) were subsampled by filtration onto 2.0 µm-pore-size Isopore filters for diatom RNA extraction (300 ml), diatom endometabolite analysis (600 ml), and pigment analysis (50 ml). RNA filters were immediately flash-frozen in liquid nitrogen and together with endometabolite analysis filters stored at -80°C until processing; pigment analysis filters were stored at -20°C.

### Growth, pigments, and photosynthetic parameters

Diatom growth was measured using a Beckman Z2 Coulter Counter, with 1 ml samples collected daily from two replicates of each temperature. All samples were diluted 10x or 100x prior to analysis on the Coulter Counter to obtain a diatom cell density in a measurable range (10^3^-10^5^ cells ml^-1^). For cell enumeration at inoculation and harvest, cells were fixed with glutaraldehyde (1% final concentration), and stored overnight at 4°C, and then at -80°C. Cells were stained with SYBR® Green (Life Technologies) and analyzed by flow cytometry (NovoCyte Quanteon, Agilent Technologies, Inc., USA). Diatom growth rates were calculated from time of inoculation until harvest as μ = ln(N_2_/N_1_)/(t_2_ − t_1_), where N is cells ml^-1^ and t is time in days. Diatom cell size at harvest was determined by Coulter Counter.

Diatom cells were analyzed for pigment composition by extracting in 2 ml 90% acetone at 4°C for 24 h. Tubes were then centrifuged at 3,000 RPM for 1.3 min (Thermo Sorvall Legend XR1), and the supernatant analyzed with a UV Vis spectrophotometer. The composition and concentration of these pigments were determined from an absorbance spectrum of 350-750 nm (54). Photosynthetic capacity (Fv/Fm) was obtained from 5 ml subsamples by analysis on a Satlantic FIRe Fluorometric System (blue light) within an hour of collection.

### Diatom endometabolite analysis

Filters for diatom endometabolite analysis were transferred into 15 ml of ultrapure MilliQ water in 50 ml tubes and sonicated in an ice bath for 7 min (50 s on, 10 s off, duty cycles 60, ∼60 µm amplitude; Branson SLPe model) to resuspend the cells. The liquid fraction was collected in new tubes and the procedure repeated three times, after which fractions were combined and stored at - 80°C until further processing. At analysis, samples were lyophilized (Labconco, Kansas City, MO, USA) and pellets mixed with 600 μl of phosphate buffer (30 mM phosphate in deuterated water, pH 7.4) and 1 mM of internal standard 2,2-dimethyl-2-silapentane-5-sulfonate. Samples were vortexed for 5 min and centrifuged at 20,800 RCF for 10 min, and supernatants were transferred to 5 mm NMR tubes (Bruker, Billerica, MA, USA). Extraction and buffer blank controls were also prepared. A pooled control sample used for annotation was prepared by combining aliquots of all samples and sample processing was carried out at 4°C.

Metabolites were analyzed by NMR spectroscopy using a 600 MHz AVANCE III HD instrument (Bruker) equipped with a 5 mm TCI cryoprobe and pulse programs of ^1^H-^13^C heteronuclear single quantum correlation (HSQC, hsqcetgpprsisp2.2 by Bruker nomenclature) and ^1^H-^13^C HSQC-total correlation spectroscopy (HSQC-TOCSY, hsqcdietgpsisp.2). TopSpin (Bruker) version 3.5 was used for NMR operation. Data were processed by NMRPipe (55). Peak intensity was extracted by rNMR version 1.11 (56) and data were analyzed by MATLAB (MathWorks) version R2023b. Peak intensity was normalized by biovolume and auto-scaled. Metabolites were annotated based on chemical shift (HSQC) and correlation information (HSQC-TOCSY). Chemical shift values for candidate peaks were obtained from the Biological Magnetic Resonance Data Bank (BRMB) (57) and the Human Metabolome Database (58), and raw reference spectra from BMRB were used for validation. Four compounds of interest that are not in these databases were annotated using literature values [homarine based on Boroujerdi et al. (59); 2,3-dihydroxypropane-1-sulfonate (DHPS), dimethylsulfoniopropionate (DMSP), and β-1,3-glucan based on Uchimiya et al. (60)]. A confidence level of annotation was assigned to each metabolite, where 1 = putative identification with functional group information; 2 = partially matched to HSQC chemical shift information in the databases or literature; 3 = fully matched to HSQC chemical shift; 4 = fully matched to HSQC chemical shift and validated by HSQC-TOCSY; 5 = validated by a spiking experiment (Table S8). All raw spectra, NMRPipe scripts for spectrum processing, and MATLAB scripts for data analysis are deposited in Metabolomics Workbench (61) under Project ID PR001837.

### Diatom mRNA analysis

RNA was extracted from filters using the ZymoBIOMICS RNA Miniprep Kit (Zymo Research, Irvine, CA, USA) according to the manufacturer’s protocol. Stranded RNAseq libraries were prepared by the Joint Genome Institute (JGI) and sequenced on a NovaSeq (Illumina). Reads were filtered and trimmed using the JGI QC pipeline, followed by evaluation of artifact sequences by kmer matching (kmer = 25) using BBDuk, allowing one mismatch; detected artifacts were trimmed from the 3’ end of the reads. Quality trimming was performed using the phred trimming method set at Q6 and reads under the minimum length of 25 bases or 1/3 of the original read length were removed. Filtered reads from each library were aligned to the reference genome using HISAT2 version 2.2.0 and strand-specific coverage bigwig files were generated using deepTools v3.1, and FeatureCounts was used to generate gene counts. *T. pseudonana* gene annotations and biosynthesis pathways were based on BioCyc (62) and PhyloDB (63).

### Metabolite lability experiment

Diatom cells harvested from the 12 axenic flasks, 4 at each temperature (Fig. 1b), were rinsed from thawed filters into 10 ml sterile L1 medium, probe sonicated on ice for 20 min to lyse cells (duty cycles 60, ∼60 µm amplitude; Branson SLPe model), and passed through pre-combusted GF/F filters. A 0.5 ml subsample of each filtered, concentrated diatom endometabolome was diluted 100-fold with fresh L1 medium and acidified with 25 µl 12 M HCl. Samples were analyzed for DOC concentration on a Shimadzu TOC-L total organic carbon analyzer coupled to a TNM-L analyzer (Mass Spectrometry Lab, Woods Hole Oceanographic Institution). Milli-Q water blanks and standard curves of potassium hydrogen phthalate and potassium nitrate were run and comparisons were made daily to standards (obtained from D. Hansell, University of Miami). To assess the lability of the compounds analyzed by NMR, a 2.7 ml aliquot of each concentrated diatom endometabolome and a blank with no endmetabolome addition were inoculated with *R. pomeroyi* (∼3 x 10^7^ cells ml^-1^) and grown for 10 h with shaking at 30°C in dark conditions. Subsamples of 1 ml were collected at inoculation and again at 10 h for metabolite quantification. From these, 540 µl were mixed with 60 µL phosphate buffer and transferred to NMR tubes, and spectra were collected using the HSQC procedure described above. One medium blank was also included. Experimental and spectrum processing settings were those described above, except TopSpin version 3.6.4 was used for NMR operation and NMRPipe was run on NMRbox (64). Endometabolite peak intensity was normalized to diatom biovolume at time of harvest.

### Data Analysis

Statistical significance of the response of diatom endometabolome composition under different acclimation temperatures was determined by the Kruskal Wallis Test with Mann Whitney U Tests for post hoc pairwise comparisons. Statistical significance of endometabolite concentrations in axenic versus co-culture endometabolomes was determined by Student’s t-tests with Benjamini-Hochberg correction for multiple hypothesis testing. Analysis for gene expression differences between *T. pseudonana* transcriptomes was carried out using DESeq2 (version 1.28.1). Sample numbers (n) were 3 or 4 for all analyses, and *P* value cutoffs were ≤ 0.05.

## Acknowledgements

The work was supported by the Simons Collaboration on Principles of Microbial Ecosystems (PriME, 542391) to MAM, the Swedish Research Council award to MO (2018–06571), and the National Science Foundation awards to MAM and AE (OCE-1948104, OCE-2019589). The work conducted by the U.S. Department of Energy Joint Genome Institute, a DOE Office of Science User Facility, is supported by the Office of Science of the U.S. Department of Energy under Contract No. DE-AC02-05CH11231 (Project ID 506891). We thank B. Hopkinson for guidance in study design and pigment analyses and the Mass Spectrometry Lab at the Woods Hole Oceanographic Institution for DOC measurements. This is the NSF Center for Chemical Currencies of a Microbial Planet (C-CoMP) publication #035.

## Author Contributions

MO, MU, ASE, and MAM conceptualized and designed the study. MO and MU conducted data curation. MAM and ASE are responsible for funding acquisition. MO, MU, FXF-G, JES, CBS, and MAP carried out investigations. MU and CBS designed methodology. MO, MU, and MAM conducted formal analysis. MO, MU, and MAM and wrote the manuscript with input from all authors.

## Data Availability Statement

Raw and annotated *T. pseudonana* transcriptome data are available through the JGI Genome Portal, proposal ID 506891 (https://genome.jgi.doe.gov/portal/) and BCO-DMO (DOI: 10.26008/1912/bco-dmo.905306.1). Metabolomics data and associated sample preparation protocols and NMR analysis and processing parameters are deposited at the Metabolomics Workbench Data Repository under Project ID PR001837 (DOI: 10.21228/M88B0T) and BCO-DMO (DOI: 10.26008/1912/bco-dmo.928203.1).

## Competing Interests

The authors declare no competing interests.

---

**Figure S1.**
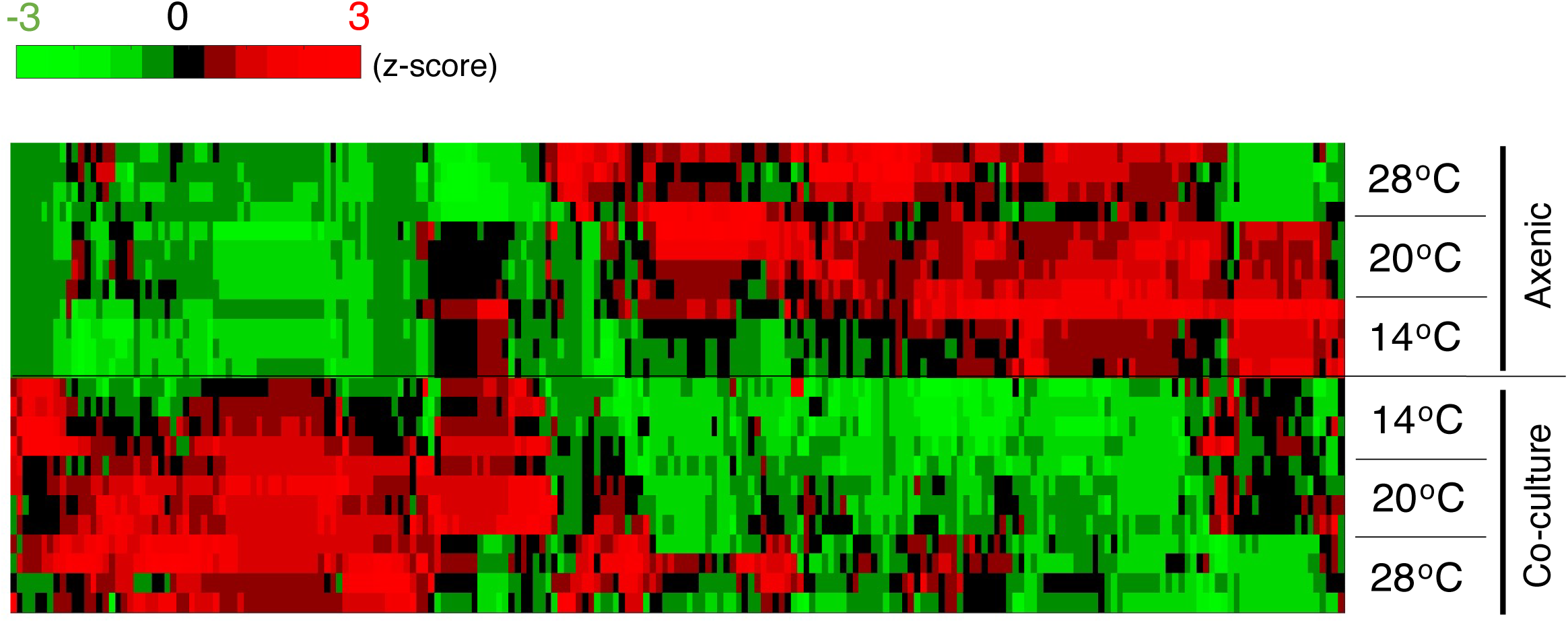
Endometabolite peak intensity from axenic (top) and co-cultured (bottom) *T. pseudonana* endometabolomes. Following acclimation at three temperatures (14°C, 20°C, and 28°C) for ∼120 generations, strains were grown to late exponential growth phase with (Co-culture) or without (Axenic) the bacterium *R. pomeroyi*. Values shown are z-scores of peak intensity following normalization to diatom biovolume at time of harvest.

**Figure S2.**
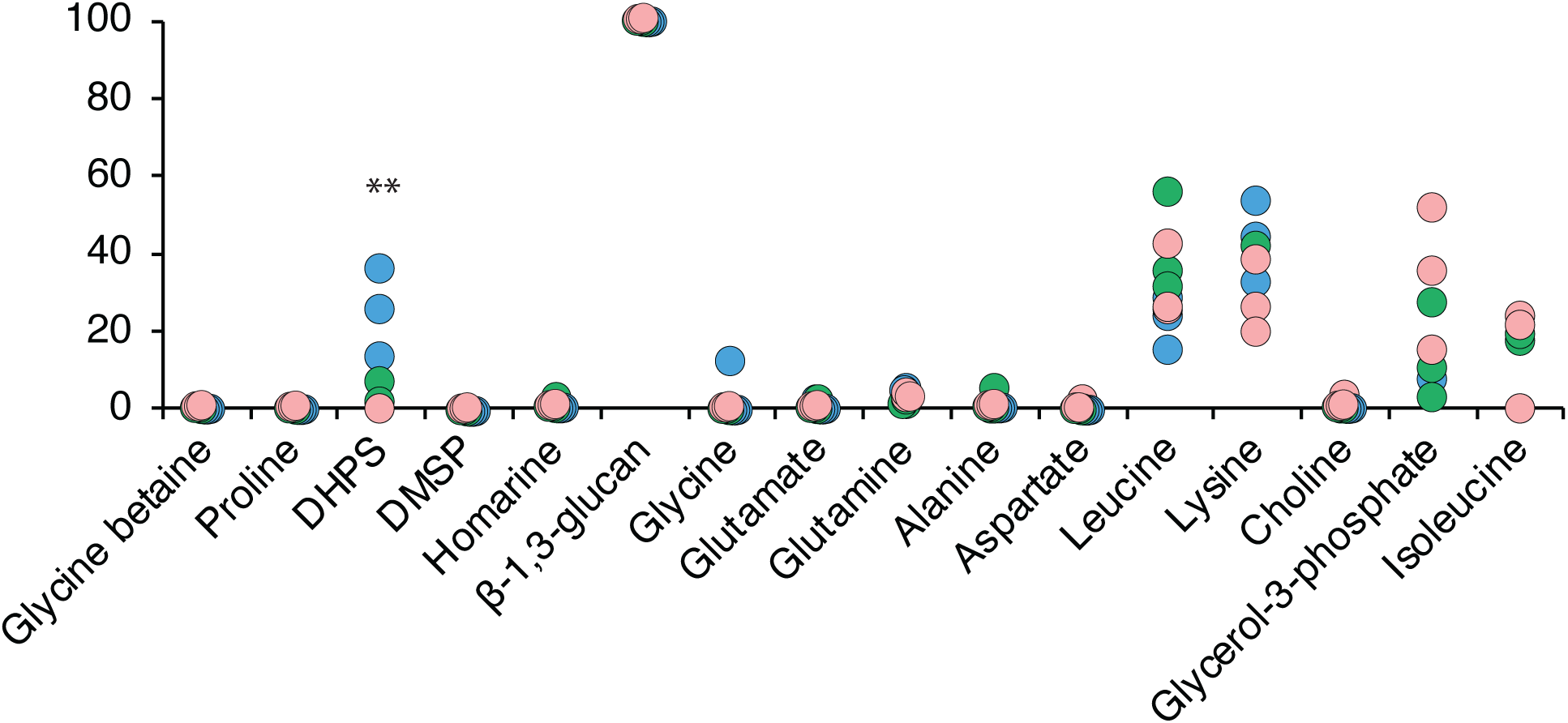
Drawdown of 16 *T. pseudonana* endometablites by *R. pomeroyi* over a 10 h incubation. Endmetabolites were extracted from diatom strains acclimated at three temperatures. Asterisks indicate a significantly lower drawdown rate of DHPS in cell lysates from diatoms acclimated at 14°C (blue) compared to 20°C (green) and 28°C (pink) (Kruskal Wallis test, p ≤ 0.01). β-1,3-glucan was not taken up by the bacterium. DHPS, 2,3-dihydroxypropane-1-sulfonate. DMSP, dimethylsulfoniopropionate.

Tables S1, S6, and S8 are provided here.

Tables S2, S3, S4, S5, and S7 are Excel files. These are available in the uploaded supplemental tables source file named “Tables S2-S5, S7”.

**Table S1.** Diatom cell physiology at harvest on day 3 (28°C treatment), day 4 (28°C), and day 6 (14°C). Chl = Chlorophyll a, Fuco = fucoxanthin, Diat = diatoxanthin, Beta-car = beta-carotene. Mean ± SD of four replicates are shown.

| **Table S1.** Diatom cell physiology at harvest on day 3 (28°C treatment), day 4 (28°C), and day 6 (14°C). Chl = Chlorophyll a, Fuco = fucoxanthin, Diat = diatoxanthin, Beta-car = beta-carotene. Mean ± SD of four replicates are shown. | | | | | | | |
| --- | --- | --- | --- | --- | --- | --- | --- |
| Category | Parameter | Axenic |  |  | Co-Culture |  |  |
|  |  | 14°C | 20°C | 28°C | 14°C | 20°C | 28°C |
| Physiology | Diameter ($\mu\text{m}$) | 3.7 ± 0.0 | 4.1 ± 0.0 | 5.4 ± 0.0 | 3.7 ± 0.0 | 4.0 ± 0.1 | 5.5 ± 0.0 |
| | Volume ($\mu\text{m}^3$) | 38.2 ± 0.0 | 54.4 ± 0.1 | 124.5 ± 3.5 | 38.2 ± 0.0 | 51.3 ± 2.1 | 130.7 ± 2.2 |
| | Abundance ($\times 10^5$ cells $\text{ml}^{-1}$) | 21.8 ± 0.7 | 17.7 ± 1.4 | 8.0 ± 0.8 | 4.5 ± 0.5 | 4.8 ± 0.3 | 4.5 ± 0.5 |
| | Biovolume ($\times 10^7$ $\mu\text{m}^3$ $\text{ml}^{-1}$) | 8.3 ± 0.3 | 9.6 ± 0.8 | 9.9 ± 0.7 | 1.7 ± 0.2 | 2.4 ± 0.2 | 6.0 ± 0.7 |
| | Fv/Fm | 0.70 ± 0.01 | 0.74 ± 0.00 | 0.72 ± 0.04 | 0.72 ± 0.01 | 0.72 ± 0.00 | 0.71 ± 0.00 |
| | Growth rate ($\text{d}^{-1}$) | 1.08 ± 0.01 | 1.60 ± 0.09 | 1.94 ± 0.17 | 0.86 ± 0.04 | 1.36 ± 0.0 | 1.71 ± 0.12 |
| Light-harvesting | Chlorophyll *a* ($\text{fg } \mu\text{m}^{-3}$) | 2.85 ± 0.08 | 4.49 ± 0.11 | 3.60 ± 0.29 | 1.46 ± 0.15 | 2.27 ± 0.17 | 2.08 ± 0.58 |
| pigments | Chlorophyll *c* ($\text{fg } \mu\text{m}^{-3}$) | 0.09 ± 0.00 | 0.19 ± 0.01 | 0.12 ± 0.02 | 0.05 ± 0.01 | 0.06 ± 0.01 | 0.07 ± 0.02 |
| | Fucoxanthin ($\text{fg } \mu\text{m}^{-3}$) | 1.08 ± 0.02 | 1.87 ± 0.10 | 1.54 ± 0.10 | 0.47 ± 0.04 | 1.01 ± 0.06 | 0.89 ± 0.25 |
| Photo-protective pigments | Beta-carotene ($\text{fg } \mu\text{m}^{-3}$) | 0.47 ± 0.02 | 0.50 ± 0.04 | 0.49 ± 0.05 | 0.30 ± 0.03 | 0.25 ± 0.05 | 0.37 ± 0.12 |
| | Diadinoxanthin ($\text{fg } \mu\text{m}^{-3}$) | 0.00 ± 0.00 | 0.00 ± 0.00 | 0.00 ± 0.00 | 0.01 ± 0.01 | 0.00 ± 0.00 | 0.00 ± 0.00 |
| | Diatoxanthin ($\text{fg } \mu\text{m}^{-3}$) | 0.19 ± 0.03 | 0.19 ± 0.04 | 0.33 ± 0.07 | 0.13 ± 0.02 | 0.29 ± 0.06 | 0.21 ± 0.01 |
| Pigment ratios |  |  |  |  |  |  |  |
| | Chl:Fuco | 2.65 ± 0.13 | 2.41 ± 0.12 | 2.34 ± 0.10 | 3.08 ± 0.18 | 2.26 ± 0.03 | 2.32 ± 0.03 |
| | Chl:Diat | 15.83 ± 3.42 | 25.48 ± 0.732 | 11.37 ± 0.36 | 11.10 ± 1.03 | 8.40 ± 2.77 | 9.66 ± 2.43 |
| | Chl:Beta-car | 6.11 ± 0.09 | 9.03 ± 0.83 | 7.41 ± 0.36 | 4.95 ± 0.11 | 9.30 ± 1.24 | 5.73 ± 0.43 |

**Table S6.**
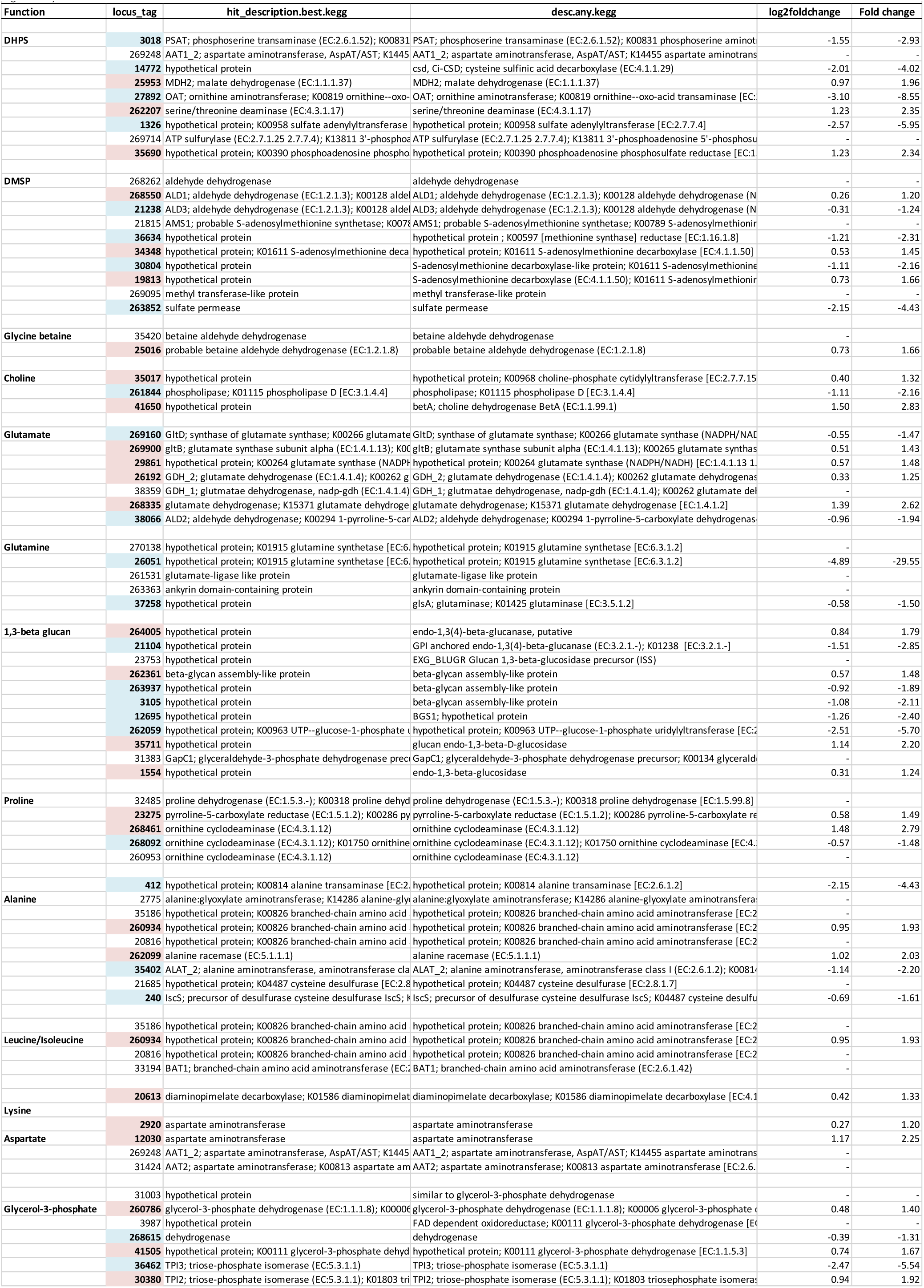
Tentatively identified Thalassiosira pseudonana metabolite synthesis genes included in Figure 4. Gene locus tags with blue shading are significantly enriched at 14°C and with pink shading are significantly enriched at 28°C.

**Table S8.** Annotation information for metabolites in this study. For the original sources of chemical shift values, see Material and Methods. Confidence level: 1 = putative identification with functional group information; 2 = partially matched to HSQC chemical shift information in the databases or literature; 3 = fully matched to HSQC chemical shift; 4 = fully matched to HSQC chemical shift and validated by HSQC-TOCSY; 5 = validated by a spiking experiment.

| Compounds | <sup>1</sup>H (ppm) | <sup>13</sup>C (ppm) | HSQC-TOCSY | Confidence level |
| --- | --- | --- | --- | --- |
| β-1,3-glucan | 3.51, 3.51, 3.56, 3.73*, 3.78°, 3.92*, 4.79 | 70.9, 78.4, 76.0, 63.4*, 86.9°, 63.4*, 105.4 | C1-C2-C3-C4-C5-C6 | 4 |
| Betaine | 3.25°, 3.89 | 55.9°, 68.6 | n.a. | 3 |
| Proline | 1.99, 2.07, 2.34, 3.32°, 3.41, 4.13 | 26.5, 31.8, 31.7, 49°, 49, 64 | 1.99-2.07/2.34, 1.99-3.32/3.414.13-2.07/2.34 | 4 |
| 2,3-dihydroxypropane-1-sulfonate (DHPS) | 3.03, 3.11°, 3.58, 3.69, 4.14 | 56.5, 56.5°, 67.3, 67.3, 70.8 | 4.14-3.03/3.11, 4.14-3.58/3.69 | 4 |
| Dimethylsulfoniopropionate (DMSP) | 2.73, 2.91°, 3.44 | 34.1, 27.8°, 43.4 | 2.73-3.44 | 4 |
| Homarine | 4.37°, 7.98, 8.04, 8.55, 8.72 | 49.3°, 130.2, 129, 149.4, 148.6 | 8.04-8.55-7.98-8.72 | 4 |
| Glutamate | 2.09*, 2.34°, 3.74* | 29.8*, 36.4°, 57.6* | 2.09-2.34/3.74 | 4 |
| Glutamine | 2.12*, 2.44°, 3.76* | 29.3*, 33.9°, 57.2* | 2.12-2.44/3.76 | 4 |
| Alanine | 1.49, 3.78° | 19.0, 53.6° | 1.49-3.78 | 4 |
| Aspartate | 2.71, 2.8, 3.91° | 39.3, 39.5, 55.1° | 3.91-2.71/2.8 | 4 |
| Leucine | 0.94°, 0.96, 1.7, 1.71*, 3.74* | 23.6°, 24.8, 42.6, 26.8*, 56.2* | 0.94/0.96-1.71, 1.7-3.74 | 4 |
| Lysine | 1.43*, 1.49*, 1.72, 1.88*, 3.02°, 3.75* | 24*, 24*, 29.2, 32.7*, 42.1°, 57.5* | 3.75-1.88-1.43/1.49-1.72-3.02 | 3 |
| Choline | 3.19*, 3.5*, 4.05° | 56.7*, 70.1*, 58.5° | 3.5-4.05 | 4 |
| Glycerol 3-phosphate | 3.61*, 3.68*, 3.78, 3.82°, 3.86* | 65*, 65*, 67.6, 67.6°, 74* | 3.86-3.61/3.68 | 3 |
| Isoleucine | 0.93, 1°, 1.25, 1.45, 1.96, 3.65* | 13.9, 17.4°, 27.2, 27, 38.7, 62.5* | – | 3 |
| *, Overlap with background peak(s). |  |  |  |  |
| a, Representative peak used for quantification. |  |  |  |  |

